# ortho2align: a sensitive approach for searching for orthologues of novel lncRNAs

**DOI:** 10.1101/2021.12.16.472946

**Authors:** Dmitry Evgenevich Mylarshchikov, Andrey Alexandrovich Mironov

**Affiliations:** Faculty of Bioengineering and Bioinformatics, Lomonosov Moscow State University, Moscow 119234, Russian Federation; Institute for Information Transmission Problems, Russian Academy of Sciences, Moscow, 127994, Russian Federation

**Keywords:** lncRNAs, evolution, orthology, software

## Abstract

**Background:** Many novel long noncoding RNAs have been discovered in recent years due to advances in high-throughput sequencing experiments. Finding orthologues of these novel lncRNAs might facilitate clarification of their functional role in living organisms. However, lncRNAs exhibit low sequence conservation, so specific methods for enhancing the signal-to-noise ratio were developed. Nevertheless, current methods such as transcriptomes comparison approaches or searches for conserved secondary structures are not applicable to novel lncRNAs dy design.

**Results:** We present ortho2align — a versatile sensitive synteny-based lncRNA orthologue search tool with statistical assessment of sequence conservation. This tool allows control of the specificity of the search process and optional annotation of found orthologues. ortho2align shows similar performance in terms of sensitivity and resource usage as the state-of-the-art method for aligning orthologous lncRNAs but also enables scientists to predict unannotated orthologous sequences for lncRNAs in question. Using ortho2align, we predicted orthologues of three distinct classes of novel human lncRNAs in six Vertebrata species to estimate their degree of conservation.

**Conclusions:** Being designed for the discovery of unannotated orthologues of novel lncRNAs in distant species, ortho2align is a versatile tool applicable to any genomic regions, especially weakly conserved ones. A small amount of input files makes ortho2align easy to use in orthology studies as a single tool or in bundle with other steps that researchers will consider sensible.

## Background

Many new long noncoding RNAs have been discovered in human cell lines in recent years due to advances in high-throughput RNA sequencing experiments [1–3]. However, some of them might come from sequencing noise or aberrations during transcript assembly. Conservation studies of novel lncRNAs might support their existence as functional genomic units and support their functional roles in cells as well as shed light on their mechanism of action [4]. But, lncRNAs exhibit low sequence conservation [5], so specific methods for enhancing the signal-to-noise ratio are needed. The first group of such methods leverages experimental evidence and compares transcriptomes of two or more species [4,6], which results in direct orthologues assignment but restricts discovery of unannotated orthologues. The second group searches for conserved secondary structures [7,8]. Still, these methods are not applicable to novel lncRNAs discovered in single original experiments, as there will be no congruent experimental data from the target species. Also, novel lncRNAs are usually not assigned with any covariational matrix and might not require any conserved secondary structure for carrying out their functions [9]. To aid the search of orthologues of novel lncRNAs we developed ortho2align — a sensitive synteny-based approach with a statistical assessment of sequence conservation. We estimated the performance of the method on a published lncRNAs orthologues dataset, compared ortho2align with a synteny-based baseline approach and a state-of-the-art transcriptome comparison approach, and predicted orthologues for 3 distinct classes of novel lncRNAs.

## Results

### ortho2align algorithm and implementation

The formal problem is as follows: to find a single orthologue in a subject genome for every lncRNA in a given annotation of the query genome. The ortho2align pipeline breaks the solution into several steps, which are described below (see also Figure 1).

**Figure 1.**
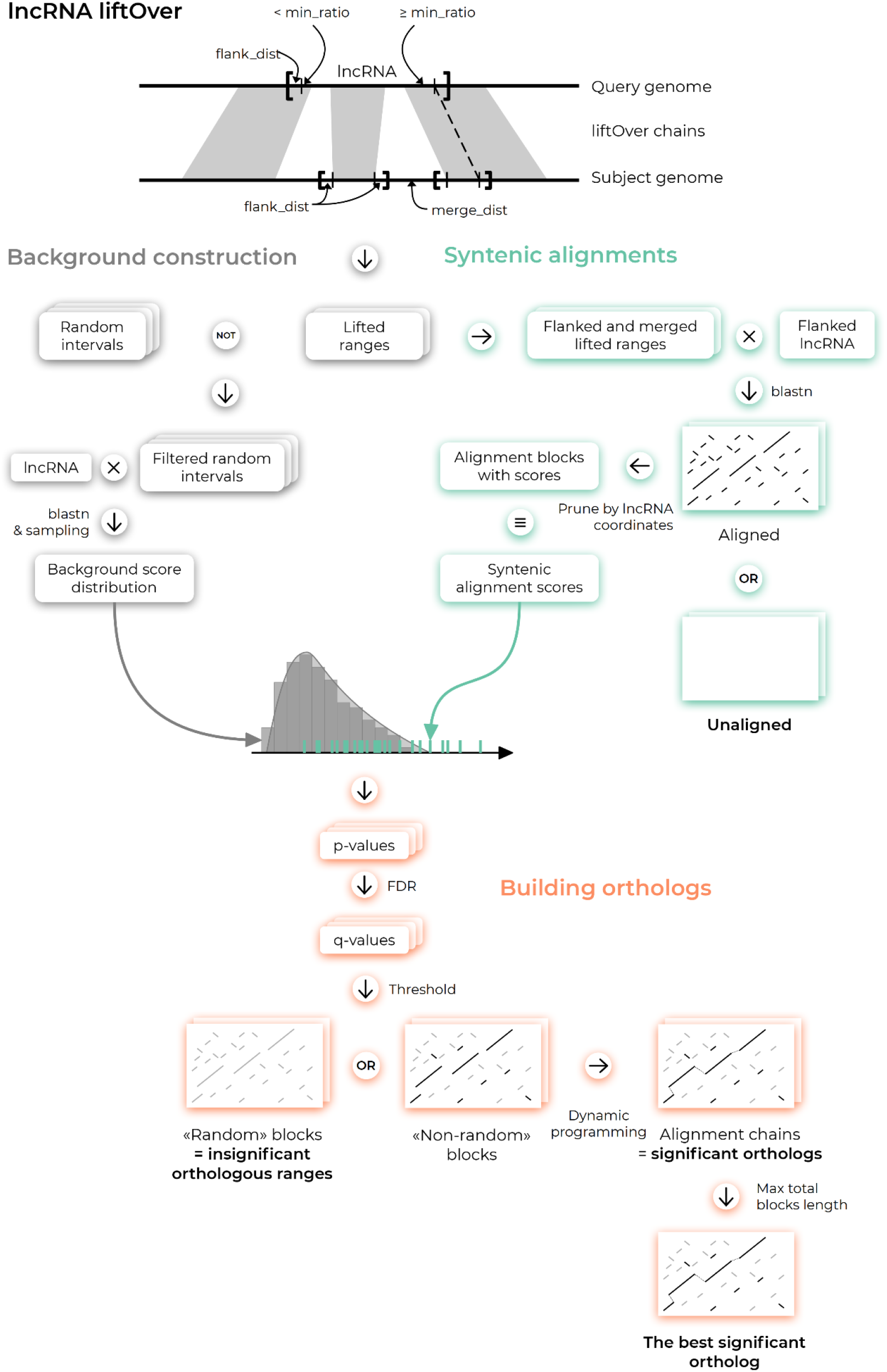
ortho2align algorithm for a single lncRNA.

#### Building syntenies

First, lncRNA coordinates are lifted from the query species genome to the subject species genome with liftOver [10] with allowed duplications and minMatch=0.05 by default. Next, syntenic regions are constructed from those lifted coordinates. Duplications are merged if they are closer than the specified distance. Constructed syntenies are flanked with a specified number of nucleotides.

#### Getting alignments

Flanked lncRNAs are aligned to their syntenic regions with BLASTN [11] with loose parameters (word_size is set to 6 by default) which results in a set of high-scoring segment pairs (HSPs) for every syntenic region for every lncRNA pruned by lncRNA coordinates. lncRNAs that were successfully lifted but have zero HSPs are deemed as *unaligned* and their lifted coordinates in subject species are reported separately.

#### Estimating background

To remove spurious HSPs a background distribution of raw HSP scores is constructed for every lncRNA by aligning its sequence to a sample of shuffled genomic ranges from the annotation of the subject genome. To ensure potential orthologues are not included in the set of background ranges, any background range is removed in case it intersects lifted coordinates of one or more lncRNAs.

#### Filtering

*HSPs*. For every lncRNA every HSP is assigned a p-value based on a right-sided statistical test in which the HSP score is so large only for random reasons based on the background distribution of raw HSP scores. P-values are adjusted with the Benjamini-Hochberg procedure. Only HSPs with a q-value less than a specified α value are retained to control the false discovery rate at α · 100% (5% by default). lncRNAs with no significant HSPs retained are deemed as *insignificant* orthologues and all their HSPs are reported separately.

#### Building orthologues

Retained HSPs are linked via dynamic programming to form alignment chains, one per every syntenic region of every lncRNA. Those chains are deemed *significant* orthologues and reported with their significant HSPs.

#### Selecting the best significant orthologue for every lncRNA

Only one orthologue for every lncRNA is selected based on the maximal sum of lengths of HSPs that comprise the orthologue in question.

#### Annotating orthologues (optional)

Found orthologues can be annotated with a provided gene annotation of the subject genome by intersecting two gene sets in a strand-specific manner. Jaccard indexes and overlap coefficients of intersecting orthologues and subject species genes are also reported.

Running BLASTN with loose parameters increases sensitivity, but applying it to the whole genome dramatically increases the working time and the number of false HSPs. Restricting the search space to syntenic regions overcomes these issues. Additionally, every lncRNA is aligned to a sample of shuffled genomic ranges of the subject genome to estimate how well it aligns to random places of the subject genome. The generated distribution of random HSP scores is used to assess whether syntenic HSPs represent truly conserved sequences or are not significantly different from random HSPs. The latter procedure enhances the specificity of the algorithm.

Besides the best significant orthologues, ortho2align reports all significant orthologues, insignificant orthologues, and lifted coordinates of unaligned lncRNAs to provide one with complete information on conservation status of as many lncRNAs as possible.

ortho2align is implemented in python 3 and requires only two non-python dependencies: standalone BLAST and liftOver programmes. Due to algorithm being atomic to a single lncRNA, ortho2align benefits much from parallelization on multiple cores. ortho2align is designed to leave a small RAM footprint by dumping intermediate files into disk so parallelization will not affect RAM usage much.

ortho2align requires only five input files: query species lncRNAs annotation, query species genome file, subject species genome file, subject species gene annotation for background ranges construction and optional annotation and a liftOver chain file (Figure S1). All these files are available from UCSC and other major annotation providers as is, so gathering them would not be a problem. Also, liftOver chain file can be generated for genome assemblies that are not stored at UCSC. As intermediate files are preserved, each step of the pipeline can be run separately (Additional file 1, Figure S1), so there is no need to run the whole pipeline again just to try different parameter values.

### Performance estimation

To estimate the performance of the method we took a public dataset of lncRNA orthologues in various species [12]. We predicted orthologues of all human lncRNAs in 6 species: rhesus macaque, mouse, opossum, platypus, chicken and *Xenopus tropicalis* — and compared the best significant predicted orthologues with true annotated ones. True positive rate (TPR) was over 80% in all species, even the distant ones, which means high sensitivity of ortho2align taking into account different evolutionary distances between human and subject species. False positive rate (FPR) dropped from 80% to 10% with increase in evolution distance between human and subject species whereas accuracy increased along the specified direction (see Figure 2). Precision was low except for macaque probably due to high class imbalance: there are much fewer human lncRNAs conserved in distant species than non-conserved as labeled in the benchmarking dataset.

**Figure 2.**
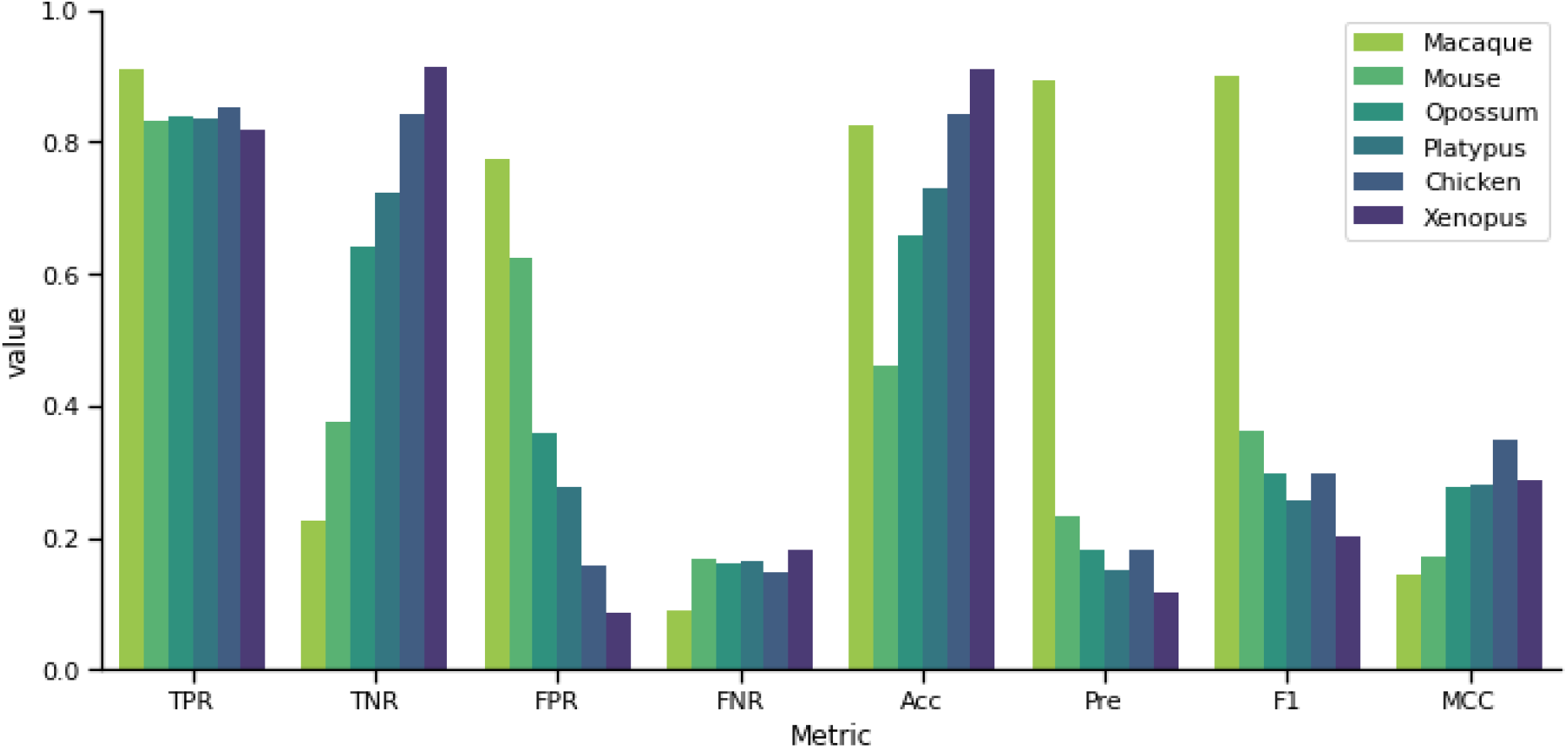
ortho2align performance metrics on the benchmarking dataset.

Taking into account all significant orthologues resulted in a small gain of TPR (up to 2.3%) but no increase in false positive predictions, which suggests an incorrect resolution of paralogues either in ortho2align or in the benchmarking dataset (Figure 3). Combining the best significant orthologues with the insignificant ones and also with unaligned lifted regions resulted in a small gain of TPR (up to 2.8%), but led to substantial increase in FPR (up to 15%). These statistics clearly state that HSPs filtering and orthologues selection steps do increase specificity and precision of the procedure.

**Figure 3.**
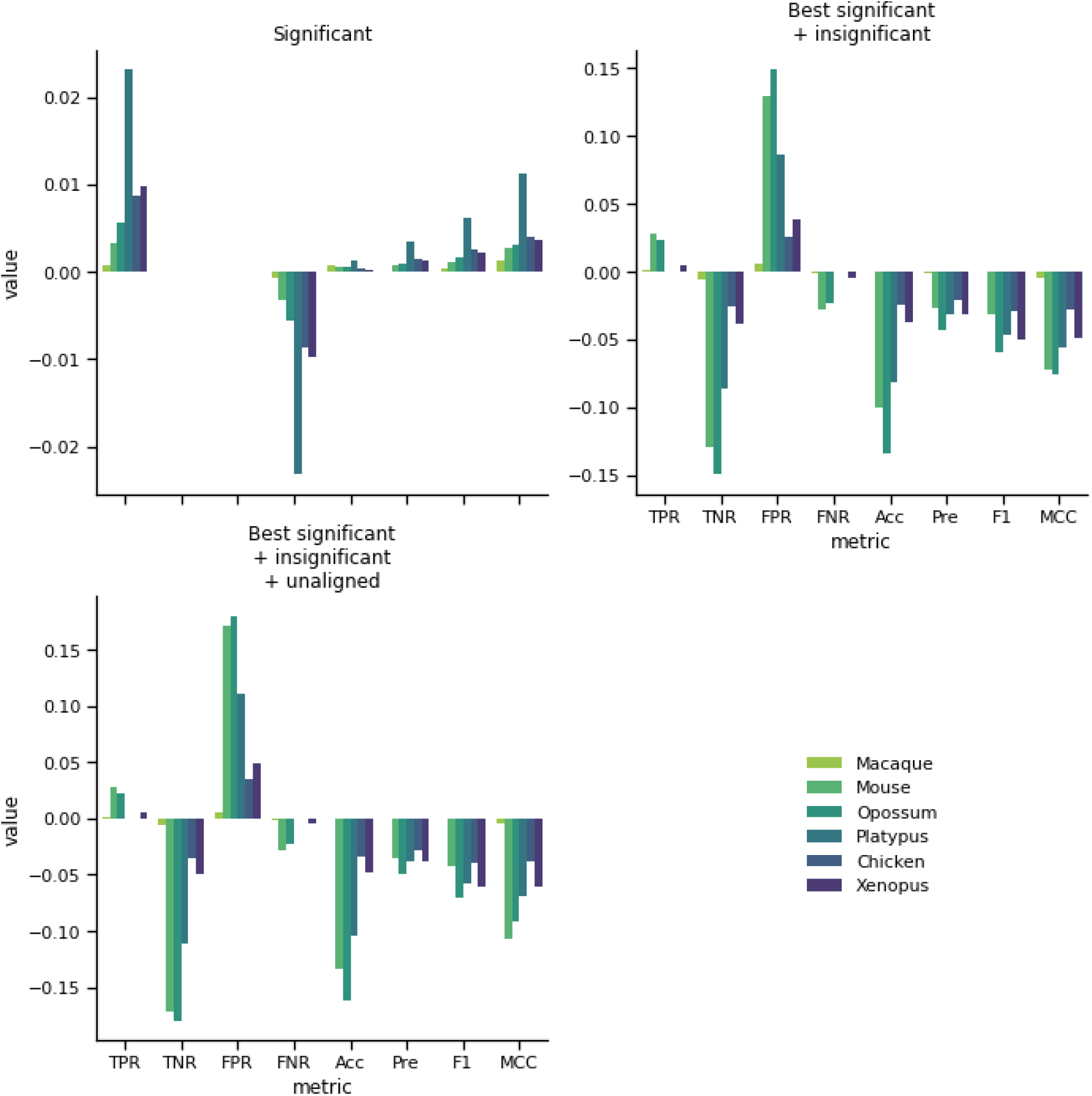
ortho2align performance metrics gain compared to the best significant orthologues compared when added other sets of predicted orthologues.

### Comparison with analogous tools

To our knowledge, there are no specific tools solving the same problem: discovery of orthologues of previously unannotated lncRNAs. So to find out how our method compares to other methods in the field, we took two analogous tools. The first one is slncky — a state-of-the art method for lncRNAs orthologue assignment based on transcriptomes comparison and restriction by synteny. The second one is a naive liftOver with allowed duplications — a baseline approach of lifting coordinates between syntenic regions. Due to slncky requiring numerous annotation files we restricted our comparison to finding orthologues of human lncRNAs in mouse as numerous files required for that were already provided by slncky.

slncky outperformed other methods in all metrics except for false negative rate (FNR), where liftOver was the best. ortho2align showed the lowest TPR, though differences between methods are quite small, but the second lowest FPR (see Figure 4). Despite not having outperforming metrics, ortho2align has two design advantages over the other tools. First, slncky searches for orthologues if the syntenic region contains one or more annotated lncRNAs, which results in a small FPR by design but also limits detection of unannotated orthologues. It might be possible that several to many orthologues have not yet been observed in any experiment, so predicting unannotated orthologues might be valuable and high FPR of ortho2align might come from those *bona fide* unannotated orthologues. Second, liftOver does not permit duplications by default, which might result in a significant loss of orthologues, or doesn’t provide a strategy to select one orthologue out of many for a single lncRNA in case duplications are allowed. ortho2align does provide this strategy, but also saves information about potential paralogues. So, ortho2align shows a comparable performance to the analogous tools in the field while being specifically crafted for the task in question: discovery of orthologues of novel lncRNAs.

**Figure 4.**
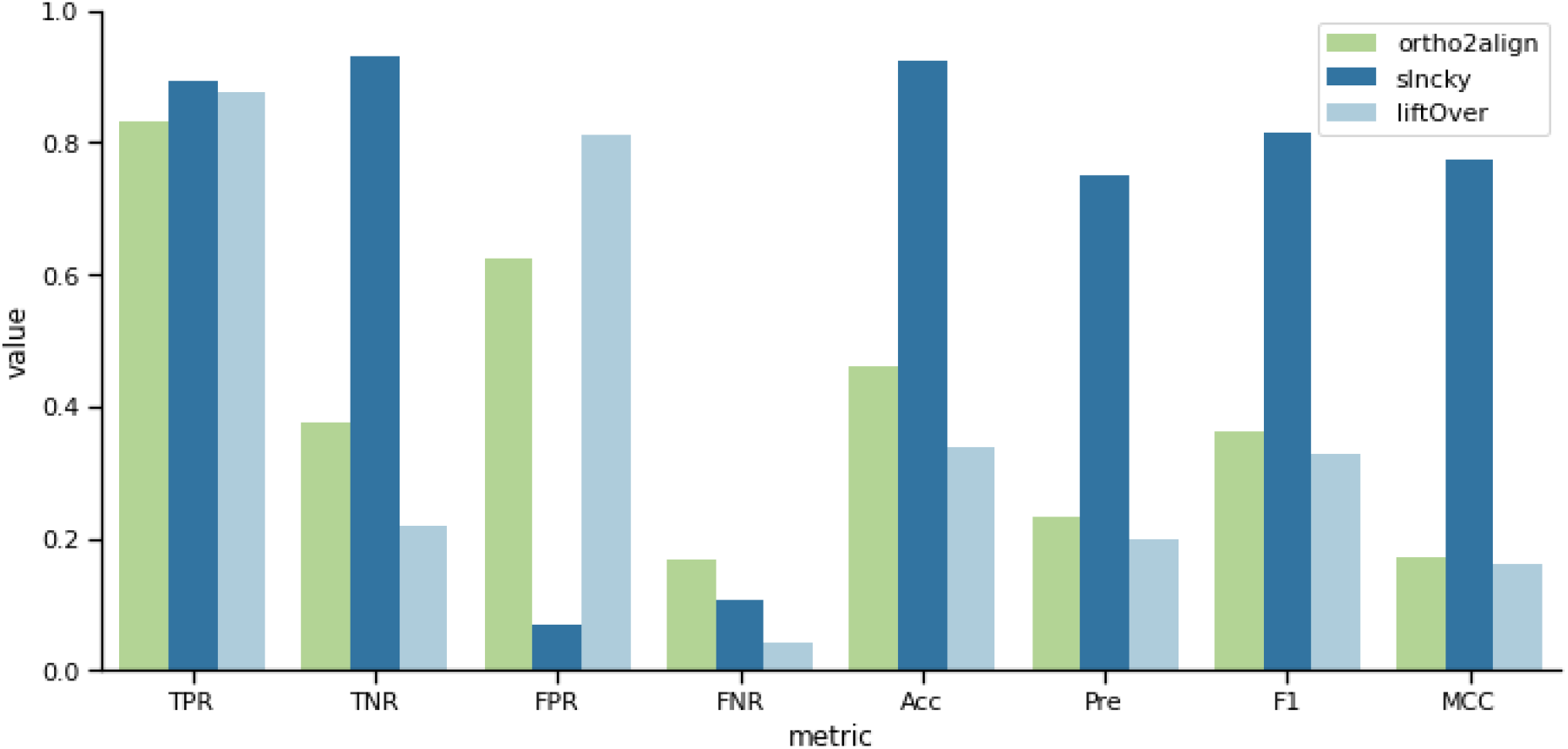
Performance metrics of ortho2align, slncky and liftOver on predicting mouse orthologues.

### Resources usage

All ortho2align, slncky and liftOver launches were done on a single node of two Intel Xeon X5670 processors. ortho2align and slncky were parallelized on 20 cores.

ortho2align running time and peak RAM usage is quite small (Table 1) due to high degree of parallelization and dumping intermediate files into disk, which makes ortho2align feasible to run on multicore machines with restricted RAM size. Resource usage of ortho2align is comparable to slncky and liftOver when predicting orthologues in mouse, which indicates no particular need in optimizing ortho2align in terms of resources.

**Table 1.** Programmes resource usage during discovery of orthologues of 14682 human lncRNAs in different species.

| Programme | Subject species | Running time | Peak RAM usage |
| --- | --- | --- | --- |
| ortho2align | Macaque | 02:43:05 | 955 Mb |
| ortho2align | Mouse | 01:42:30 | 1 Gb 749 Mb |
| slncky | Mouse | 01:31:39 | 1 Gb 563 Mb |
| liftOver | Mouse | 00:00:59 | 1 Gb 565 Mb |
| ortho2align | Opossum | 02:23:59 | 789 Mb |
| ortho2align | Platypus | 01:19:16 | 534 Mb |
| ortho2align | Chicken | 00:47:07 | 247 Mb |
| ortho2align | Xenopus | 01:01:11 | 208 Mb |

### Discovery of orthologues of novel human lncRNAs

To present practical usage and significance of our approach we aimed to predict orthologues of three distinct classes of novel human lncRNAs: novel chromatin-associated RNAs (X-RNAs) were discovered in a Red-C experiment, are enriched in active or repressed chromatin and may modulate chromatin structure [3], short tandem repeat-enriched RNAs (strRNAs) might interact with multiple copies of specific RNA-binding proteins [2] and semi-extractable RNAs (seRNAs) are thought to comprise novel nuclear bodies [1].

X-RNAs exhibited a certain decrease in the number of significant orthologues with increase of the evolutionary distance from human to the subject species (see Figure 5). 210 X-RNAs out of 1865 were significantly conserved in all six species and 625 X-RNAs were shown to be primate-specific. Most X-RNAs weakened their conservation statuses with increase of the evolutionary distance with a few exceptions.

**Figure 5.**
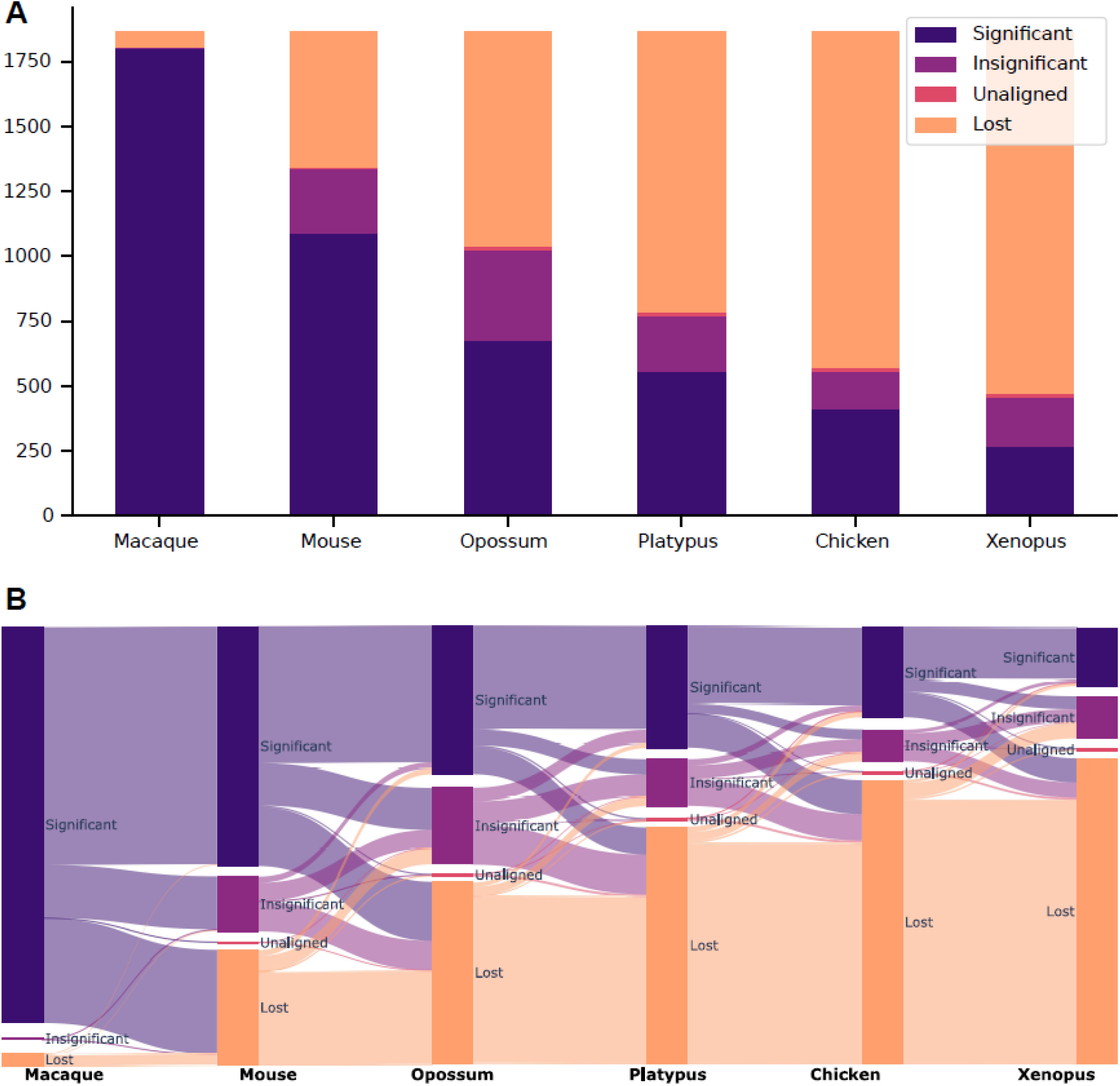
Predicted orthologues of X-RNAs in six Vertebrata species. A. Distribution of conservation statuses across species. B. Conservation status flow between species.

In accordance with X-RNAs being defined as intergenic transcripts in a strand-specific manner, most of their orthologues were also intergenic with no regard to species or conservation status (Additional file 1, Figure S2).

strRNAs were found to be much less conserved with no significantly conserved RNAs across all species and 34 out of 96 RNAs being primate-specific (Additional file 1, Figure S3). Notably, PNCTR, one of strRNAs and the architectural RNA of the perinucleolar compartment, was found to be significantly conserved in all but two species, Opossum and Platypus, where it was insignificantly conserved, but successfully aligned to the respective syntenic regions.

Concerning three novel seRNAs, only one of them, peri-NPIP RNA, was significantly conserved in all six species (Additional file 1, Figure S4). PZP antisense RNA was found to be significantly conserved in all species but Mouse, where it was insignificantly conserved. SVA D RNA has shown a stable decrease in its conservation with a total loss in Platypus and Chicken (probably due to being located at X chromosome in Human), yet unexpectedly was syntenic to a certain region in the *X. tropicalis* genome.

Annotations of lncRNAs orthologues are available in Additional file 2.

## Discussion

We developed ortho2align — a synteny-based approach for finding orthologues of novel lncRNAs with a statistical assessment of sequence conservation. Implemented strategies of restricting the search to syntenic regions, statistical filtering of HSPs and selection of orthologues provide high levels of sensitivity and specificity as well as optimal computational time even when looking for orthologues in distant species. ortho2align has shown a little bit lower TPR compared to two analogous tools, which is probably due to either incorrect paralogue resolution in the benchmarking dataset or more stringent statistical HSP filtering in ortho2align pipeline compared to slncky: the latter does not apply a multiple testing correction procedure during filtering, which is statistically inaccurate. Although analogous tools outperform ortho2align a little in terms of quality metrics, they are either restricted to a narrow case of the comparison of two lncRNAs annotations derived from transcriptomic data or too primitive to provide a researcher with essential information on the conservation of given lncRNAs. On the contrary, being designed for the discovery of unannotated orthologues of novel lncRNAs in distant species, ortho2align is a versatile tool applicable to any genomic regions, especially weakly conserved ones. Researchers are also provided with full information about loci conservation status: either there is only syntenic relationship, two loci are alignable or their alignments are statistically significant. ortho2align also retains information about putative paralogues along with the best orthologues selected out of them. This versatility, small amount of input files and the completeness of information complemented with optional annotation of orthologues will allow researchers to adapt ortho2align in their orthology studies. ortho2align can also be used in bundle with other steps that researchers will consider sensible, such as the best reciprocal hit strategy [13] or removal of possible protein-coding orthologues [6].

To display the predictive power of ortho2align, we predicted orthologues of three distinctive classes of novel human lncRNAs. Multiple X-RNAs were found to be conserved even in *X. tropicalis*, and their orthologues were mostly intergenic as X-RNAs themselves, which in total hints functional significance of those novel chromatin-associated RNAs. Annotation of murine X-RNAs in the RADICL [14] or GRID [15] data will allow for direct inference of human-to-mouse orthologous X-RNAs and support the supposition of their functional significance. The high degree of conservation of PNCTR, one of the most prominent strRNAs, indicates its significant role in maintaining the perinucleolar compartment in mammalian cells. Conservation of other strRNAs also suggests their putative significance in binding multiple molecules of the same protein. Finally, the analysis of conserved seRNAs might lead to the discovery of novel nuclear bodies with those seRNAs as their architectural components. Application of ortho2align to other classes of novel lncRNAs will likely be as fruitful.

## Conclusions

We developed ortho2align — a synteny-based approach for finding orthologues of novel lncRNAs with a statistical assessment of sequence conservation. The approach is sensitive and specific considering its ability to discover unannotated orthologues. These features allowed us to predict orthologues for three distinct classes of novel human lncRNAs. The versatility and low resource usage make ortho2align suitable for a wide range of applications in orthology studies. ortho2align source code, list of dependencies, default parameter values, installation and testing instructions are publically available at https://github.com/dmitrymyl/ortho2align under GPL-3.0 license.

## Methods

### Dataset of orthologues

We used a publicly available dataset of lncRNA orthologues in various species [12]. We took lncRNAs from the Supplementary dataset 1 of the aforementioned publication. We chose lncRNAs annotations of the main dataset for 7 species: human, rhesus macaque, mouse, opossum, platypus, chicken and Xenopus tropicalis. Due to liftOver files are being readily available only for UCSC assemblies and dataset lncRNAs were annotated in accordance with Ensembl assemblies, we mapped Ensembl genome versions to UCSC genome versions (Additional file 1, Table S1) and manually converted chromosome names from Ensembl to UCSC nomenclature with some lncRNAs in contigs being lost (Additional file 1, Table S2). We then prepared tables mapping names of orthologous lncRNAs in human and every other species from one-to-one lncRNA families data. Annotation and mapping data is available in Additional file 3.

### Quality metrics definition

A true positive hit is defined as a predicted orthologue intersecting a correct orthologue in a strand-specific manner. A false positive hit is defined as an orthologue predicted for a lncRNA with no annotated orthologues in the dataset. TP is defined as a number of true positive hits, FP is defined as a number of false positive hits, TN is defined as a total number of non-orthologous lncRNAs (total_non_ortho) minus FP and FN is defined as a total number of orthologous lncRNAs (total_ortho) minus TP. Ratio metrics are defined as follows: TPR = TP / total_ortho, FPR = FP / total_non_ortho, TNR = TN / total_non_ortho, FNR = FN / total_ortho, accuracy = (TP + TN) / (TP + TN + FP + FN), precision = TP / (TP + FP), F1 = TP / (TP + 0.5 * (FP + FN)).

Due to duplications, liftOver might find multiple orthologues for a single lncRNA, so its TP and FP values are weighted after calculating TN and FN. Every lncRNA contributes not a “1” to TP or FP, but a share of a true (or false) orthologue(-s) to the found orthologues.

### Benchmarking settings

ortho2align v1.0.4 was run with run_pipeline subcommand. Default parameter values were used except for min_ratio, pval_threshold, and FDR correction. The best min_ratio value was selected between 0.01 and 0.05. The best pval_threshold value was selected among the following values: 1e-6, 1e-2, 5e-2, 1e-1, 2e-1. Selection of the best values of those two parameters was conducted jointly by maximizing TPR for each species separately (see Additional file 1, Table S3 for the final values). FDR correction was enabled. Background genomic ranges were constructed from the subject species lncRNAs annotation files from the benchmarking dataset. The same annotation files were used for the annotation of found orthologues. The programme was parallelized on 20 cores. Running time and RAM consumption was recorded by the programme itself.

slncky v1.0.4 was run with key -2 (no filtering, only searching for orthologues) and default parameter values. Annotation and genome files that were used were provided with slncky except for the annotation of noncoding RNAs in mm9, which was replaced with mm9 lncRNAs annotation from the benchmarking dataset. The programme was parallelized on 20 cores. Running time and RAM consumption was recorded with the programme “GNU time 1.7”.

liftOver v377 was run with the following parameters: -multiple -noSerial -minMatch=0.05 (a default minMatch value in the UCSC online utility). Predicted orthologues were annotated with bedtools intersect -s -loj -a <predicted_orthologues> -b <true_orthologues>, where <true_orthologues> is the mm9 lncRNAs annotation from the benchmarking dataset. Running time and RAM consumption was recorded with the programme “GNU time 1.7”. Genome fasta files and liftOver chain files for respective genome versions were downloaded from the UCSC website and were used in ortho2align and liftOver.

### Discovery of orthologues of novel human lncRNAs

We have downloaded annotations of X-RNAs, strRNAs and seRNAs from the respective publications. We aimed at predicting their orthologues in the most recent genomes assemblies of species (Additional file 1, Table S4), yet corresponding liftOver chain files were available only for hg38 assembly. As X-RNAs and seRNAs were annotated for hg19 assembly, we converted them to hg38 assembly with liftOver (only two out of 1867 X-RNAs and no seRNAs were lost). strRNAs were already annotated for hg38 assembly. We ran ortho2align with parameters selected during the benchmarking process. To annotate predicted orthologues we used RefSeq annotations for the corresponding genomes (Additional file 1, Table S4) with chromosome names converted between RefSeq and UCSC systems using UCSC chromAlias files and a custom script. Only genes (i.e. records with gbkey = “Gene”) were used for the annotation process.

## List of abbreviations

HSP: high-scoring segment pair
TP: true positive
FP: false positive
TN: true negative
FN: false negative
TPR: true positive rate
FPR: false positive rate
FNR: false negative rate
TNR: true negative rate
Acc: accuracy
Pre: precision
F1: F1-score
X-RNAs: novel chromatin-associated RNAs
strRNAs: short tandem repeat-enriched RNAs
seRNAs: semi-extractable RNAs

## Declarations

### Ethics approval and consent to participate

Not applicable.

### Consent for publication

Not applicable.

### Availability of data and materials

All data generated or analysed during this study are included in this published article and its supplementary information files.

### Authors’ contributions

AM proposed the idea of the research, DM developed the algorithm, obtained the results described in the manuscript and wrote the manuscript, AM revised and edited the manuscript. All authors read and approved the final manuscript.

## Acknowledgements

Not applicable.

