## Additional file 1 for "ortho2align: a sensitive approach for searching for orthologues of novel lncRNAs"

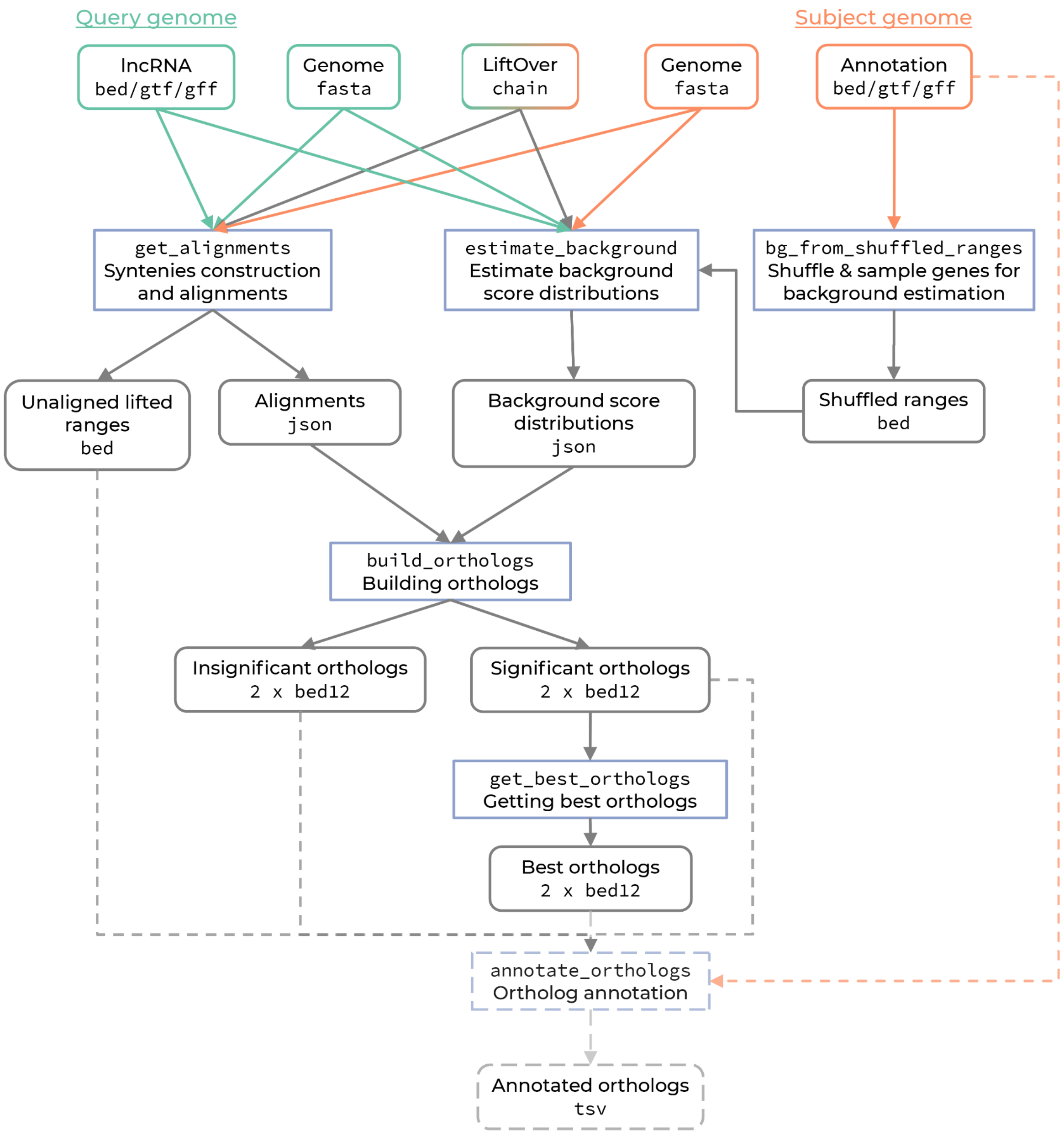


**Figure S1.** ortho2align pipeline composition.

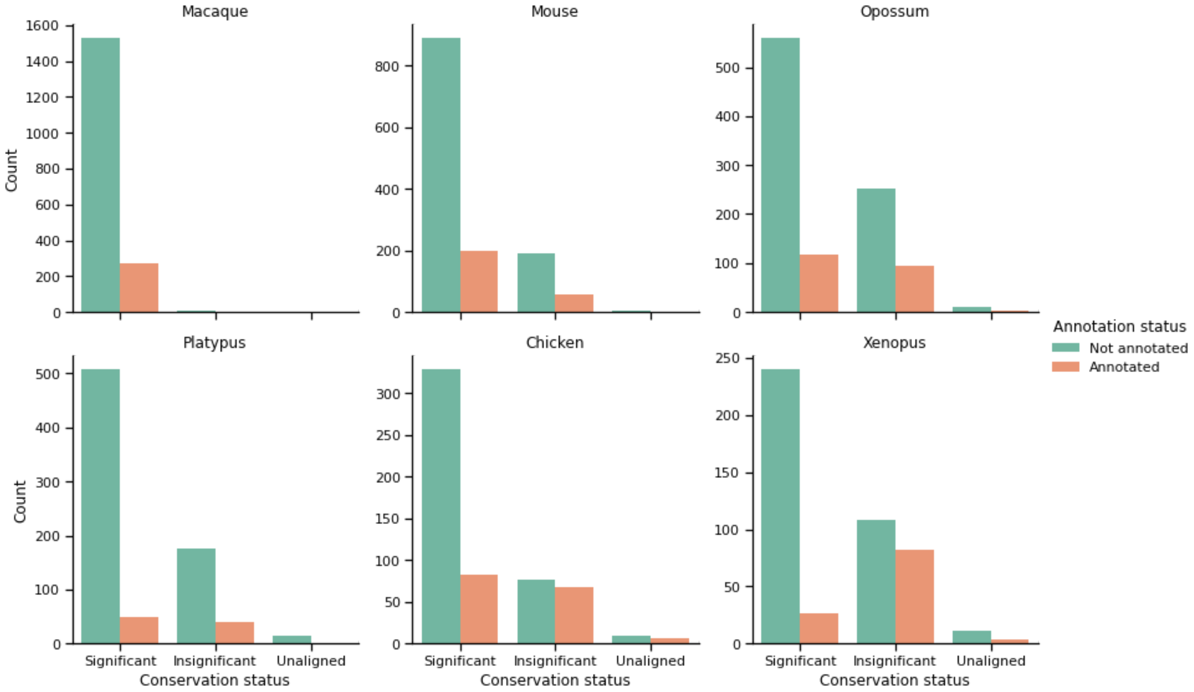


**Figure S2.** Annotation status of X-RNAs orthologues across species and conservation statuses.

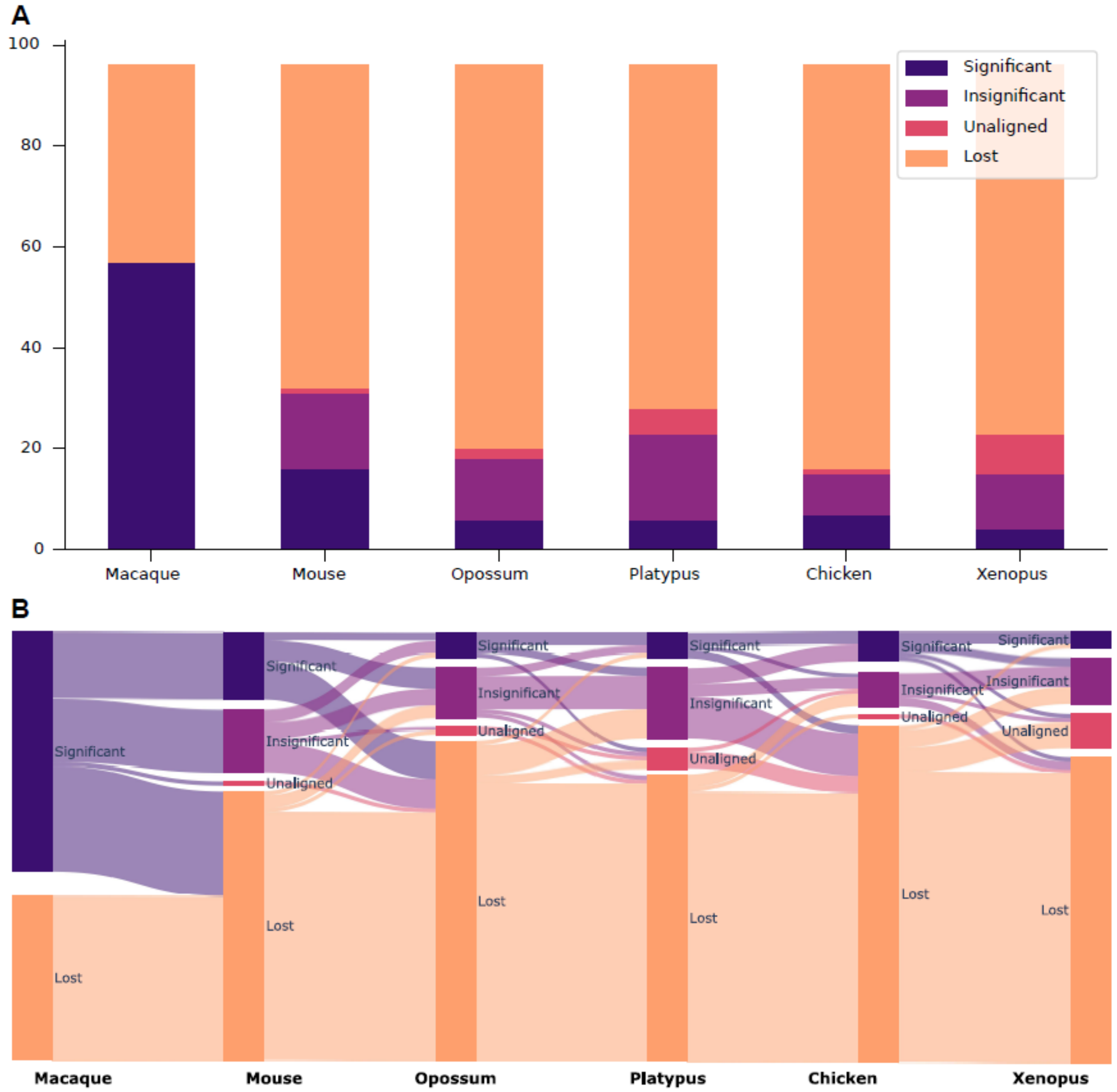


**Figure S3.** Predicted orthologues of strRNAs in six Vertebrata species.
**A.** Distribution of conservation statuses across species. **B.** Conservation status flow between species.

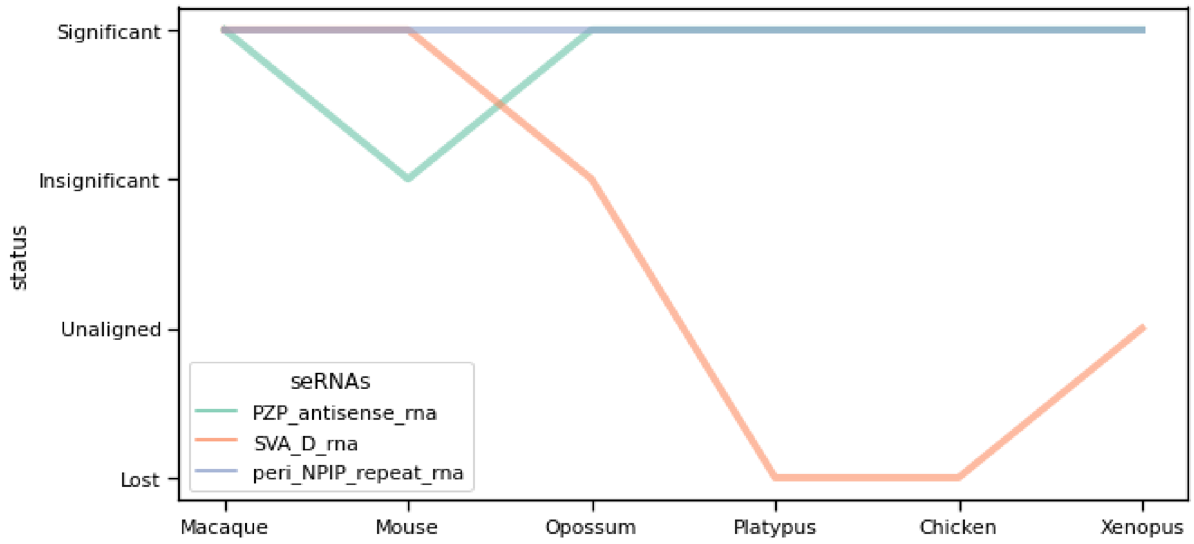


**Figure S4.** Unannotated seRNAs orthologues' conservation statuses.

**Table S1.** Genome versions used in benchmarking.

| Species | Ensembl genome version | UCSC genome version | Assembly level |
|---|---|---|---|
| Human | GRCh37 | hg19 | Chromosome |
| Macaque | MMUL 1.0 | rheMac2 | Chromosome |
| Mouse | NCBI37 | mm9 | Chromosome |
| Opossum | monDom5 | monDom5 | Chromosome |
| Platypus | OANA5 | ornAna1 | Chromosome (78% in scaffolds) |
| Chicken | WASHUC2 | galGal3 | Chromosome |
| Xenopus | JGI 4.2 | xenTro3 | Scaffold |

**Table S2.** lncRNAs orthologues dataset statistics.

| Species | lncRNAs, total | lncRNAs lost in chromosome names conversion | lncRNAs retained in chromosome names conversion | Orthologues of human lncRNAs |
|---|---|---|---|---|
| Human | 14682 | 0 | 14682 | Not applicable |
| Macaque | 15280 | 559 (from contigs) | 14721 | 12868 |
| Mouse | 10850 | 9 | 10841 | 2720 |
| Opossum | 8039 | 0 | 8039 | 1261 |
| Platypus | 6889 | 32 (from contigs) | 6857 | 823 |
| Chicken | 5412 | 0 | 5412 | 580 |
| Xenopus | 3296 | 0 | 3296 | 204 |

**Table S3.** Best parameter values of ortho2align in terms of TPR maximization.

| Species | min_ratio | pval_threshold |
|---|---|---|
| Macaque | 0.05 | 0.2 |
| Mouse | 0.01 | 0.2 |
| Opossum | 0.01 | 0.1 |
| Platypus | 0.01 | 0.05 |
| Chicken | 0.01 | 0.1 |
| Xenopus | 0.01 | 1e-6 |

**Table S4.** Genome versions used in predicting orthologues for novel human lncRNAs.

| Species | UCSC version | RefSeq version |
|---|---|---|
| Macaque | rheMac10 | GCF_003339765.1 |
| Mouse | mm10 | GCF_000001635.26 |
| Opossum | monDom5 | GCF_000002295.2 |
| Platypus | ornAna2 | GCF_000002275.2 |
| Chicken | galGal6 | GCF_000002315.5 |
| Xenopus | xenTro10 | GCF_000004195.4 |
