## Additional file 2 for "ortho2align: a sensitive approach for searching for orthologues of novel lncRNAs": stats.txt

-----------------------
estimate_background stats:
Recieved 3 transcripts.
Eliminated 0 background ranges due to intersections with lifted transcripts.
Estimated background for 3 of them.
Caught 0 exceptions.
Distribution of amount of found background HSPs:
2| 5,
-----
2| 61,
-----
10| 15,
Reported at most 1000 HSPs for each transcript.
-----------------------
-----------------------
get_alignments stats:
Recieved 3 transcripts.
Aligned 2 of them.
Estimated as unliftable 1 transcripts.
Estimated as unaligned 0 transcripts.
Caught 0 exceptions.
Distribution of amount of alignments:
1: 2
Reported all alignments for each transcript.
-----------------------
-----------------------
build_orthologs stats:
Recieved 2 transcript alignments and 2 backgrounds.
Built orthologs for 2 of transcripts.
Found only unsignificant orthologs for 0 transcripts.
Caught 0 exceptions.
Distribution of amount of orthologs:
1: 2
Reported all significant orthologs for each transcript.
-----------------------
-----------------------
annotate_orthologs stats:
Recieved 41 orthologs.-----------------------
-----------------------
annotate_orthologs stats:
Recieved 55 orthologs.-----------------------
-----------------------
annotate_orthologs stats:
Recieved 2 orthologs from galGal6/build_files/significant.subject_orthologs.bed.
Distribution of amount of annotations:
0: 1
4: 1
Reported all annotations for each ortholog.
-----------------------
-----------------------
annotate_orthologs stats:
Recieved 2 orthologs from galGal6/bestSignificant.subject_orthologs.bed.
Distribution of amount of annotations:
0: 1
4: 1
Reported all annotations for each ortholog.
-----------------------
Elapsed time: 00:01:11.
Maximum RAM usage: 0 Gb 315 Mb.
