## Additional file 2 for "ortho2align: a sensitive approach for searching for orthologues of novel lncRNAs": stats.txt

-----------------------
estimate_background stats:
Recieved 3 transcripts.
Eliminated 0 background ranges due to intersections with lifted transcripts.
Estimated background for 3 of them.
Caught 0 exceptions.
Distribution of amount of found background HSPs:
9| 5,
-----
15| 51,
16| 79,
Reported at most 1000 HSPs for each transcript.
-----------------------
-----------------------
get_alignments stats:
Recieved 3 transcripts.
Aligned 3 of them.
Estimated as unliftable 0 transcripts.
Estimated as unaligned 0 transcripts.
Caught 0 exceptions.
Distribution of amount of alignments:
1: 1
3: 1
7: 1
Reported all alignments for each transcript.
-----------------------
-----------------------
build_orthologs stats:
Recieved 3 transcript alignments and 3 backgrounds.
Built orthologs for 2 of transcripts.
Found only unsignificant orthologs for 1 transcripts.
Caught 0 exceptions.
Distribution of amount of orthologs:
0: 1
2: 1
3: 1
Reported all significant orthologs for each transcript.
-----------------------
-----------------------
annotate_orthologs stats:
Recieved 41 orthologs.-----------------------
-----------------------
annotate_orthologs stats:
Recieved 1 orthologs from monDom5/build_files/insignificant.subject_orthologs.bed.
Distribution of amount of annotations:
1: 1
Reported all annotations for each ortholog.
-----------------------
-----------------------
annotate_orthologs stats:
Recieved 5 orthologs from monDom5/build_files/significant.subject_orthologs.bed.
Distribution of amount of annotations:
0: 3
1: 2
Reported all annotations for each ortholog.
-----------------------
-----------------------
annotate_orthologs stats:
Recieved 2 orthologs from monDom5/bestSignificant.subject_orthologs.bed.
Distribution of amount of annotations:
1: 2
Reported all annotations for each ortholog.
-----------------------
Elapsed time: 00:01:42.
Maximum RAM usage: 0 Gb 767 Mb.
