## Additional file 2 for "ortho2align: a sensitive approach for searching for orthologues of novel lncRNAs": stats.txt

-----------------------
estimate_background stats:
Recieved 96 transcripts.
Eliminated 0 background ranges due to intersections with lifted transcripts.
Estimated background for 96 of them.
Caught 0 exceptions.
Distribution of amount of found background HSPs:
0| 0,0,0,0,0,0,0,0,0,0,0,0,0,0,0,0,0,0,0,0,0,0,0,0,0,0,1,1,1,2,3,3,4,5,6,7,8,
1| 2,3,6,7,7,9,
2| 0,0,0,0,3,4,5,5,6,7,8,9,
3| 6,9,
4| 3,4,5,
5| 0,1,4,7,
7| 3,9,
-----
1| 05,06,10,
2| 07,59,
3| 91,
4| 00,55,64,
6| 31,34,34,61,
7| 05,11,76,
8| 00,40,61,
9| 41,93,
-----
10| 20,92,
11| 44,
15| 50,
16| 20,
18| 56,
21| 66,
22| 37,
26| 58,
Reported at most 1000 HSPs for each transcript.
-----------------------
-----------------------
get_alignments stats:
Recieved 96 transcripts.
Aligned 32 of them.
Estimated as unliftable 64 transcripts.
Estimated as unaligned 1 transcripts.
Caught 0 exceptions.
Distribution of amount of alignments:
0: 1
1: 27
2: 4
Reported all alignments for each transcript.
-----------------------
-----------------------
build_orthologs stats:
Recieved 31 transcript alignments and 31 backgrounds.
Built orthologs for 16 of transcripts.
Found only unsignificant orthologs for 15 transcripts.
Caught 0 exceptions.
Distribution of amount of orthologs:
0: 15
1: 16
Reported all significant orthologs for each transcript.
-----------------------
-----------------------
annotate_orthologs stats:
Recieved 1 orthologs from mm10/align_files/subject_unaligned.bed.
Distribution of amount of annotations:
0: 1
Reported all annotations for each ortholog.
-----------------------
-----------------------
annotate_orthologs stats:
Recieved 18 orthologs from mm10/build_files/insignificant.subject_orthologs.bed.
Distribution of amount of annotations:
0: 12
1: 6
Reported all annotations for each ortholog.
-----------------------
-----------------------
annotate_orthologs stats:
Recieved 16 orthologs from mm10/build_files/significant.subject_orthologs.bed.
Distribution of amount of annotations:
0: 12
1: 3
2: 1
Reported all annotations for each ortholog.
-----------------------
-----------------------
annotate_orthologs stats:
Recieved 16 orthologs from mm10/bestSignificant.subject_orthologs.bed.
Distribution of amount of annotations:
0: 12
1: 3
2: 1
Reported all annotations for each ortholog.
-----------------------
Elapsed time: 00:00:39.
Maximum RAM usage: 1 Gb 585 Mb.
