## Additional file 2 for "ortho2align: a sensitive approach for searching for orthologues of novel lncRNAs": stats.txt

-----------------------
estimate_background stats:
Recieved 96 transcripts.
Eliminated 0 background ranges due to intersections with lifted transcripts.
Estimated background for 96 of them.
Caught 0 exceptions.
Distribution of amount of found background HSPs:
0| 0,0,0,0,0,0,0,0,0,0,0,0,0,0,0,0,0,0,0,0,0,0,0,0,0,0,0,1,1,1,2,2,2,2,2,2,2,3,5,5,5,6,6,7,7,
1| 2,2,3,4,7,7,7,8,8,9,
2| 0,0,2,3,5,6,
3| 1,2,2,4,4,4,5,6,6,9,
4| 1,2,4,4,6,6,9,
5| 3,6,9,
7| 4,6,
8| 0,5,
9| 3,
-----
1| 00,04,
2| 76,
7| 73,
-----
11| 38,
12| 95,
14| 59,
15| 51,
18| 26,
27| 78,
Reported at most 1000 HSPs for each transcript.
-----------------------
-----------------------
get_alignments stats:
Recieved 96 transcripts.
Aligned 28 of them.
Estimated as unliftable 68 transcripts.
Estimated as unaligned 5 transcripts.
Caught 0 exceptions.
Distribution of amount of alignments:
0: 5
1: 10
2: 5
3: 4
4: 2
5: 1
6: 1
Reported all alignments for each transcript.
-----------------------
-----------------------
build_orthologs stats:
Recieved 23 transcript alignments and 23 backgrounds.
Built orthologs for 6 of transcripts.
Found only unsignificant orthologs for 17 transcripts.
Caught 0 exceptions.
Distribution of amount of orthologs:
0: 17
1: 6
Reported all significant orthologs for each transcript.
-----------------------
-----------------------
annotate_orthologs stats:
Recieved 8 orthologs from ornAna2/align_files/subject_unaligned.bed.
Distribution of amount of annotations:
0: 7
1: 1
Reported all annotations for each ortholog.
-----------------------
-----------------------
annotate_orthologs stats:
Recieved 46 orthologs from ornAna2/build_files/insignificant.subject_orthologs.bed.
Distribution of amount of annotations:
0: 38
1: 8
Reported all annotations for each ortholog.
-----------------------
-----------------------
annotate_orthologs stats:
Recieved 6 orthologs from ornAna2/build_files/significant.subject_orthologs.bed.
Distribution of amount of annotations:
0: 6
Reported all annotations for each ortholog.
-----------------------
-----------------------
annotate_orthologs stats:
Recieved 6 orthologs from ornAna2/bestSignificant.subject_orthologs.bed.
Distribution of amount of annotations:
0: 6
Reported all annotations for each ortholog.
-----------------------
Elapsed time: 00:00:32.
Maximum RAM usage: 0 Gb 588 Mb.
