## Additional file 2 for "ortho2align: a sensitive approach for searching for orthologues of novel lncRNAs": stats.txt

-----------------------
estimate_background stats:
Recieved 96 transcripts.
Eliminated 0 background ranges due to intersections with lifted transcripts.
Estimated background for 96 of them.
Caught 0 exceptions.
Distribution of amount of found background HSPs:
0| 0,0,0,0,0,0,0,0,0,0,0,0,0,0,0,0,0,0,0,0,0,0,0,0,0,0,1,1,1,1,2,2,2,4,4,5,5,5,5,6,9,9,9,
1| 0,0,0,1,2,5,9,
2| 1,2,4,5,7,8,
3| 4,6,
4| 1,
5| 7,8,9,
6| 4,
9| 4,
-----
1| 06,09,42,56,57,65,
4| 11,
-----
11| 09,
17| 80,
52| 82,
57| 01,
62| 96,
65| 38,
66| 60,
72| 68,
76| 66,
90| 88,
96| 68,
-----
103| 18,
106| 86,
107| 47,
113| 80,80,
121| 31,
124| 70,
125| 22,
127| 79,
130| 53,
212| 90,
245| 83,
299| 27,
387| 57,
Reported at most 1000 HSPs for each transcript.
-----------------------
-----------------------
get_alignments stats:
Recieved 96 transcripts.
Aligned 57 of them.
Estimated as unliftable 39 transcripts.
Estimated as unaligned 0 transcripts.
Caught 0 exceptions.
Distribution of amount of alignments:
1: 50
2: 3
3: 3
4: 1
Reported all alignments for each transcript.
-----------------------
-----------------------
build_orthologs stats:
Recieved 57 transcript alignments and 57 backgrounds.
Built orthologs for 57 of transcripts.
Found only unsignificant orthologs for 0 transcripts.
Caught 0 exceptions.
Distribution of amount of orthologs:
1: 50
2: 3
3: 3
4: 1
Reported all significant orthologs for each transcript.
-----------------------
-----------------------
annotate_orthologs stats:
Recieved 42 orthologs.-----------------------
-----------------------
annotate_orthologs stats:
Recieved 56 orthologs.-----------------------
-----------------------
annotate_orthologs stats:
Recieved 69 orthologs from rheMac10/build_files/significant.subject_orthologs.bed.
Distribution of amount of annotations:
0: 48
1: 20
2: 1
Reported all annotations for each ortholog.
-----------------------
-----------------------
annotate_orthologs stats:
Recieved 57 orthologs from rheMac10/bestSignificant.subject_orthologs.bed.
Distribution of amount of annotations:
0: 39
1: 17
2: 1
Reported all annotations for each ortholog.
-----------------------
Elapsed time: 00:01:18.
Maximum RAM usage: 0 Gb 823 Mb.
