## Additional file 2 for "ortho2align: a sensitive approach for searching for orthologues of novel lncRNAs": stats.txt

-----------------------
estimate_background stats:
Recieved 96 transcripts.
Eliminated 1 background ranges due to intersections with lifted transcripts.
Estimated background for 96 of them.
Caught 0 exceptions.
Distribution of amount of found background HSPs:
0| 0,0,0,0,0,0,0,0,0,0,0,0,0,0,0,0,0,0,0,0,0,0,0,0,0,0,0,0,0,1,1,1,1,2,2,3,3,3,4,4,4,5,5,5,6,6,6,7,8,9,
1| 0,0,0,1,1,2,2,3,4,4,5,8,8,
2| 0,0,1,3,4,7,7,8,9,
3| 4,5,7,7,7,
4| 1,9,
5| 3,5,6,7,
6| 0,1,8,
7| 2,2,4,5,6,
8| 1,2,
-----
1| 26,
2| 16,72,
Reported at most 1000 HSPs for each transcript.
-----------------------
-----------------------
get_alignments stats:
Recieved 96 transcripts.
Aligned 23 of them.
Estimated as unliftable 73 transcripts.
Estimated as unaligned 8 transcripts.
Caught 0 exceptions.
Distribution of amount of alignments:
0: 8
1: 7
2: 2
3: 2
4: 1
5: 1
6: 1
7: 1
Reported all alignments for each transcript.
-----------------------
-----------------------
build_orthologs stats:
Recieved 15 transcript alignments and 15 backgrounds.
Built orthologs for 4 of transcripts.
Found only unsignificant orthologs for 11 transcripts.
Caught 0 exceptions.
Distribution of amount of orthologs:
0: 11
1: 4
Reported all significant orthologs for each transcript.
-----------------------
-----------------------
annotate_orthologs stats:
Recieved 13 orthologs from xenTro10/align_files/subject_unaligned.bed.
Distribution of amount of annotations:
0: 8
1: 5
Reported all annotations for each ortholog.
-----------------------
-----------------------
annotate_orthologs stats:
Recieved 30 orthologs from xenTro10/build_files/insignificant.subject_orthologs.bed.
Distribution of amount of annotations:
0: 25
1: 5
Reported all annotations for each ortholog.
-----------------------
-----------------------
annotate_orthologs stats:
Recieved 4 orthologs from xenTro10/build_files/significant.subject_orthologs.bed.
Distribution of amount of annotations:
0: 4
Reported all annotations for each ortholog.
-----------------------
-----------------------
annotate_orthologs stats:
Recieved 4 orthologs from xenTro10/bestSignificant.subject_orthologs.bed.
Distribution of amount of annotations:
0: 4
Reported all annotations for each ortholog.
-----------------------
Elapsed time: 00:00:27.
Maximum RAM usage: 0 Gb 309 Mb.
